# A kinetic-aware approach to infer metabolic variations and flux using transcriptomics and metabolomics data

**DOI:** 10.64898/2026.09.27.754711

**Authors:** Haiqi Zhu, Changlin Wan, Min Yang, Yue Fang, Zheng An, Paveethran Swaminathan, Pengtao Dang, Zhi Li, Jia Wang, Yijie Wang, Yabing Chen, Anjun Ma, Qin Ma, Mark R. Kelley, Sha Cao, Melissa L. Fishel, Chi Zhang

**Affiliations:** Department of Computer Science, Luddy School of Informatics, Computing, and Engineering, Indiana University Bloomington, Bloomington, IN 47408, USA; Department of Biomedical Engineering, Center for Biomedical Data Sciences, Knight. Cancer Institute, and Brenden-Colson Center for Pancreatic Care, Oregon Health & Science University, Portland, OR 97201, USA; Department of Pathology and Laboratory Medicine, School of Medicine, Oregon Health & Science University, Portland, OR 97239, USA; Department of Biomedical Informatics, College of Medicine, The Ohio State University, Columbus, OH 43210, USA; Department of Pediatrics and Herman B Wells Center for Pediatric Research, Indiana University School of Medicine, Indianapolis, IN 46202, USA; Indiana University Melvin and Bren Simon Comprehensive Cancer Center, Indiana University School of Medicine, Indianapolis, IN 46202, USA; Department of Biochemistry, Molecular Biology and Pharmacology, Indiana University School of Medicine, Indianapolis, IN 46202, USA

**Author notes:** **Correspondence:** Chi Zhang, Melissa Fishel, Sha Cao.

## Abstract

Assessing metabolic variations and flux quantities enable systematic understandings of metabolic shifts, reprogramming, adaptation and interactions in human diseases. However, omics-based estimation of metabolic flux and its variation remains challenging due to several fundamental limitations: the need for disease and tissue context specific metabolic model; nonlinear enzyme kinetic model that links enzyme and substrate changes to reaction flux; partial, unpaired, and snap-shot measurements across omics modalities; and uncertainty in computational prediction. Here, we present Michaelis–Menten model-based Flux Estimation Analysis (mmFEA), a Monte Carlo framework for estimating condition-specific flux changes by integrating paired or unpaired metabolomics and transcriptomics (or proteomics) data. mmFEA separates each reaction-rate change into enzyme- and substrate-associated components and assess reaction rate using Michaelis-Menten kinetics equation. A baseline metabolite saturation rate is introduced by integrating protein language model predicted kinetic parameters and human baseline level metabolic concentration to enable kinetic-aware integration of unpaired substrate and enzyme level measurements. Distribution of metabolic flux and variations between conditions are further computed using MCMC sampling by treating Michaelis-Menten-derived marginal flux distribution as prior and coherency in flux balance as likelihood. To benchmark mmFEA, we generated an in house multi-omics data set including transcriptomics, metabolomics, metabolic activity functional assay, and CRISPR screening data using pancreatic cancer cell line system treated by APEX1 inhibitors. We demonstrated that mmFEA could accurately capture experimentally observed metabolic changes and achieved a better performance than all baseline methods. Our analysis revealed the necessity in using both substrate and enzyme modality and kinetic aware model in metabolic flux assessment. Further analysis using independent pancreatic cancer cohorts further validated the robustness of mmFEA, supporting integration of condition-linked unpaired data. Pan-cancer and spatial multi-omic applications demonstrated the use of mmFEA for resolving context-dependent metabolic variation when direct flux measurements are unavailable. Together, mmFEA provides a mechanistically grounded framework for estimating relative metabolic flux changes and their uncertainty from heterogeneous omics data.

## INTRODUCTION

Metabolic fluxes quantify the rates at which biochemical reactions convert metabolites into energy, redox equivalents and biosynthetic products, thereby providing a functional description of cellular and tissue-level metabolism.[1] Metabolic reprogramming is therefore defined by altered reaction fluxes rather than by changes in metabolite abundance or enzyme expression alone. Flux alterations are characteristic of cancer, inflammatory disorders and neurodegenerative diseases and can reveal disease mechanisms, metabolic vulnerabilities and responses to genetic or therapeutic perturbations. [21 [31 [41] However, most molecular studies of human disease or experimental systems measure static abundances of transcripts, proteins or metabolites rather than the biochemical reaction rates that connect them. This limitation is particularly consequential in heterogeneous tissues, where metabolic states can vary across cell types and tissue sites. Although changes in gene expression and metabolite abundance provide information about metabolic regulation, neither modality alone directly specifies flux: enzyme abundance reflects potential catalytic capacity, whereas metabolite abundance reflects the balance of production, consumption and transport. Inferring changes in biochemical reaction rates from static molecular measurements therefore remains a fundamental challenge in studying metabolism in human disease.

Stable-isotope tracing and metabolic flux analysis can directly quantify selected reaction rates in experimentally controlled systems, but their requirements for tracer administration, isotope-resolved measurements and context-specific metabolic models limit their scalability to large clinical cohorts, archived specimens and spatially resolved human tissues.[5].[6] Emerging spatial metabolomics, such as imaging mass spectrometry, can characterize the tissue distribution of metabolites but generally do not directly determine the biochemical reaction rates responsible for those distributions.[7] Consequently, transcriptomic, proteomic, and metabolomic profiling remain the most widely available resources for studying disease-associated metabolism. Inferring dynamic flux changes from such static observations is fundamentally underdetermined: multiple metabolic states may be consistent with the same measured molecular abundances.jfil A computational framework must therefore incorporate biochemical knowledge and network-level constraints while explicitly representing uncertainty in the inferred flux states.

A growing number of constraint-based, statistical-learning, and artificial-intelligence approaches have been developed to infer metabolic activity from omics data. [8] [9] [10] These methods have substantially expanded the scale at which metabolic states can be analyzed, including single-cell and spatial resolution **(Table 1** and **Supplementary Table 1**). However, many existing approaches rely primarily on transcriptomic data and consequently approximate enzymatic capacity rather than the realized rates of metabolic reactions. Metabolomics provides complementary information about substrate availability, but metabolite abundance cannot be interpreted as flux without considering the enzymes that catalyze the reaction and the kinetic state of the system. In particular, the same change in substrate abundance may strongly affect an unsaturated reaction but have little effect when the corresponding enzyme is already saturated.[11] These observations motivate an explicitly multimodal formulation in which transcriptomic or proteomic measurements characterize changes in enzymatic capacity, metabolomics characterizes changes in substrate availability, and enzyme kinetics determines how these changes jointly affect reaction rates. Such integration must also accommodate practical limitations of existing datasets, including unmatched transcriptomic and metabolomic cohorts, relative rather than absolute metabolite measurements, incomplete metabolomic coverage, and missing kinetic parameters.

**Table 1.** Existing methods and challenges, method comparison with mmFEA.

| Approach | Input data | Model and output | Flux-balance constraint | Enzyme kinetics | In-silico Perturbation | Principal limitation |
| --- | --- | --- | --- | --- | --- | --- |
| **13C-MFA** [6] | Isotope tracing, MS/NMR isotopologues, exchange rates | Isotopomer modeling; experimentally calibrated fluxes | Stoichiometric and isotope balance | No | No | Experimentally intensive; limited scalability to clinical and spatial samples |
| **COMPASS** [12] | scRNA-seq or tissue transcriptomics data | Expression-weighted FBA; relative reaction scores | Strict steady-state balance | No | No | Does not incorporate measured metabolites or quantify posterior uncertainty |
| **scFEA** [13] | scRNA-seq or bulk transcriptomics data | Graph neural network; sample- or cell-level relative fluxes capacity | Soft balance constraint | No | Partial | Substrate availability and kinetic saturation are not modeled |
| **METAFlux** [14] | scRNA-seq or bulk transcriptomics data | Reaction activity scoring and quadratic-programming FBA | Strict steady-state balance | No | No | Depends on expression, nutrient and objective assumptions |
| **REMI** [15] | Relative transcriptomics, metabolomics, and thermodynamics | Thermodynamically constrained FBA | Strict steady-state balance | No | No | Molecular changes enter as relative constraints rather than kinetic rate laws |
| **INTEGRATE** [16] | Transcriptomics and intracellular metabolomics | Flux sampling plus reaction-expression and substrate-propensity scores | Strict steady-state balance | Mass-action approximation | No | Primarily classifies regulatory contributions rather than estimating kinetic flux distributions |
| **Metabolic digital twin** [17] | Patient-specific 13C tracing, tumor/blood measurements and scRNA-seq | Stoichiometric/isotopic simulations and neural networks; patient- and cell-type-specific fluxes | Stoichiometric and isotope conservation; supports non-steady-state settings | Isotope dynamics; no explicit enzyme-substrate kinetics | Partial | Requires specialized, patient-specific isotope-tracing data |
| **mmFEA** | Paired or unpaired transcriptomics/proteomics, metabolomics, Km information, metabolic network and reference flux | Michaelis-Menten-informed probabilistic inference; posterior flux changes and condition-specific flux distributions | Soft, bounded imbalance, supports non-steady-state settings | Michaelis-Menten-informed enzyme-substrate kinetic | Michaelis-Menten equation-based simulation | Depends on reference-flux calibration, enzyme-abundance proxies and kinetic-data coverage |

The need for mechanistically grounded metabolic flux modeling is also important with the emergence of virtual-cell and virtual-tissue models.[18] Recent artificial-intelligence models can learn cellular representations and predict molecular responses to genetic or pharmacological perturbations.[19] However, many such predictions are characterized primarily by gene expression or latent-embedding spaces. Although these representations can capture associations between perturbations and cellular states, changes in an embedding or transcriptional profile do not directly specify how the rates of biochemical reactions are altered. Mechanistically explicit in silico perturbation analysis requires a model that connects changes in enzymes and substrates to reaction-level dynamics while accounting for metabolic-network structure. Because omics data provide only partial observations of the metabolic system, the resulting constraints typically define a feasible flux solution space rather than uniquely identifying the underlying flux state. Probabilistic inference and sampling are therefore needed to characterize uncertainty across this space, quantify confidence in predicted flux changes and distinguish robust perturbation responses from changes that remain weakly constrained by the available data.

To address these challenges, we developed Michaelis-Menten model-based Flux Estimation Analysis (mmFEA), a probabilistic framework for inferring condition-specific metabolic flux distributions and flux changes through the integration of transcriptomic or proteomic data, metabolomics data, enzyme kinetic parameters, and metabolic-network structure. mmFEA first establishes a distribution of reference flux states using a compatible network-constrained flux estimator. It then applies Michaelis–Menten kinetics to decompose the fold change of each reaction rate into an enzyme-associated component and one or more substrate-associated components, thereby explicitly accounting for the effect of baseline substrate saturation. By modeling condition-specific molecular measurements as probability distributions, mmFEA can integrate paired or unpaired transcriptomic and metabolomic datasets and propagate measurement and model uncertainty into reaction-level flux estimates. Finally, mmFEA uses network-level sampling to obtain posterior flux distributions that remain biochemically coherent while allowing disease or perturbed states to deviate from strict steady-state assumptions. This formulation enables the estimation of relative flux changes and condition-specific flux magnitudes, provides confidence intervals for individual reactions, and attributes each predicted alteration to enzyme, substrate, kinetics, and network-level contributions. mmFEA can be applied independently or integrated with existing flux-estimation methods that provide reference flux distributions.

We evaluated mmFEA through complementary experimental and public-data benchmarks. We first analyzed condition-linked transcriptomic, metabolomic, and mitochondrial functional data under controlled metabolic perturbations, enabling assessment of agreement between predicted flux changes and measured functional responses. We also evaluated robustness using datasets collected in different batches and under related but nonidentical experimental contexts. A separate Supplementary analysis used CRISPR screening data to explore candidate metabolic dependencies rather than to validate the Pa03C flux estimates. Finally, we applied mmFEA to paired and unpaired transcriptomic and metabolomic datasets across cancer types and to paired spatial multi-omics data, evaluating its ability to reconstruct disease- and tissue-site-specific metabolic alterations. Together, these analyses evaluate mmFEA as an uncertainty-aware and mechanistically interpretable framework for inferring relative metabolic flux changes from heterogeneous omics data.

## RESULTS

### Mathematical considerations

To connect static molecular measurements to changes in metabolic reaction rates, we represented a metabolic network as a directed factor graph in which reactions are modeled as variables and metabolites connect reactions through their substrate-product relationships. For a reaction *R* catalyzed by enzyme *E* and involving substrate *S*, denoted as 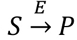, the Michaelis-Menten equation detailed below explicitly models how its reaction rate depends on the concentration of enzymes and substrates:[24]

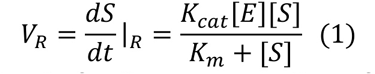

where[*E*], [*S*], *K*_*cat*_, and *K*_*m*_ stands for the concentration of enzyme *E* and substrate *S*, and reaction and kinetic parameters, respectively.

By equation (1), changes in enzymes and substrates both affect the reaction rate. Thus, a kinetic-informed flux estimator needs to leverage changes in both modalities under a correct systems biology formulation. The main challenges for developing such a capability include (1) differential equation-based fitting of Michaelis-Menton model requires knowing both kinetic parameters and measuring exact concentration of enzymes and substrates, (2) the disparities in demanded expertise and experimental systems make transcriptomics (or proteomics) and metabolomics data always collected from different batches, and (3) the omics data collected from disease samples are always non-time courses and in relative abundance level.

To address the challenges, rather than directly attempting to estimate *V*_*R*_ from incomplete molecular measurements, mmFEA estimates the fold change in reaction rate between a perturbed condition and a reference condition, denoted as *t* and c, respectively. Under the assumption that the catalytic turnover of a given reaction remains constant between conditions, *K*_*cat*_, cancels from the ratio, yielding

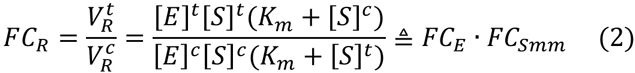

where 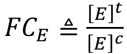 represents the enzyme-associated component, and 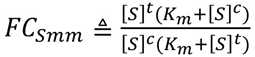 represents the substrate-associated component of flux change. Thus, the formula (2) separates transcriptomic or proteomic measurements and metabolomics measurements in characterizing changes in reaction rate.

To make the kinetic effect of substrate abundance interpretable, we defined the Baseline Saturation Ratio (BSR) for substrate Sin reaction *R* as 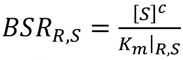, then *FC*_*S*_ can be written as

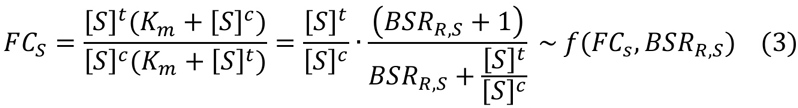

which is a function of the substrate fold change of 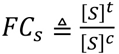 and *BSR*_*R*_,_*s*_*·* With this transformation and pre-computed BSR for all reactions and substrates, the fold change of flux rate can be written as

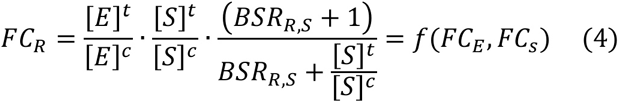

in which *FC*_*E*_ can be estimated using transcriptomic or proteomic while *FC*_*s*_ can be estimated using metabolomic data. The formula (4) form the mathematical foundation of mmFEA. Using (4), the marginal distribution of *FC*_*R*_ for each reaction can be computed using transcriptomics (or proteomics) and metabolomics and fluxome distribution of each condition can be further computed by minimizing their flux imbalance and maximizing the coherency between the marginal *FC*_*R*_ and the *FC*_*R*_ in the computed fluxome. Noted, as the term *FCE* and *FC*_*5*_ are disjoint in (4), the method enables the integration of unpaired data.

### Overall framework of Michaelis – Menten model-based Flux Estimation Analysis (mmFEA)

mmFEA is designed as a kinetic-aware metabolic flux analysis framework that integrates transcriptomic or proteomic data, metabolomic data, enzyme kinetic information, and metabolic network structure to infer condition-specific flux distributions and flux changes between conditions. **Figure 1a-g** illustrates the computational framework of mmFEA. The regular input of mmFEA include transcriptomic or proteomic and metabolomic data of two conditions — one experimental and one reference condition, while its minimal input is one modality of two conditions or fold change **(Figure 1a)**. Users can select to use mmFEA encoded metabolic networks or provide customized network (see details in Supplementary Methods). mmFEA accepts relative mass spectrometry intensities such as mass spectrometry (MS)- or NMR-based metabolomics, or absolute metabolite concentrations (quantitative metabolomics), and can analyze multi-modality generated from the same sample (paired data) or from independent studies (unpaired data). Rather than requiring every molecular feature to be observed, mmFEA represents measured, predicted, and missing quantities as probability distributions, allowing uncertainty arising from incomplete molecular coverage and kinetic information to be retained throughout the analysis **(Figure 1b**).

**Figure 1.**
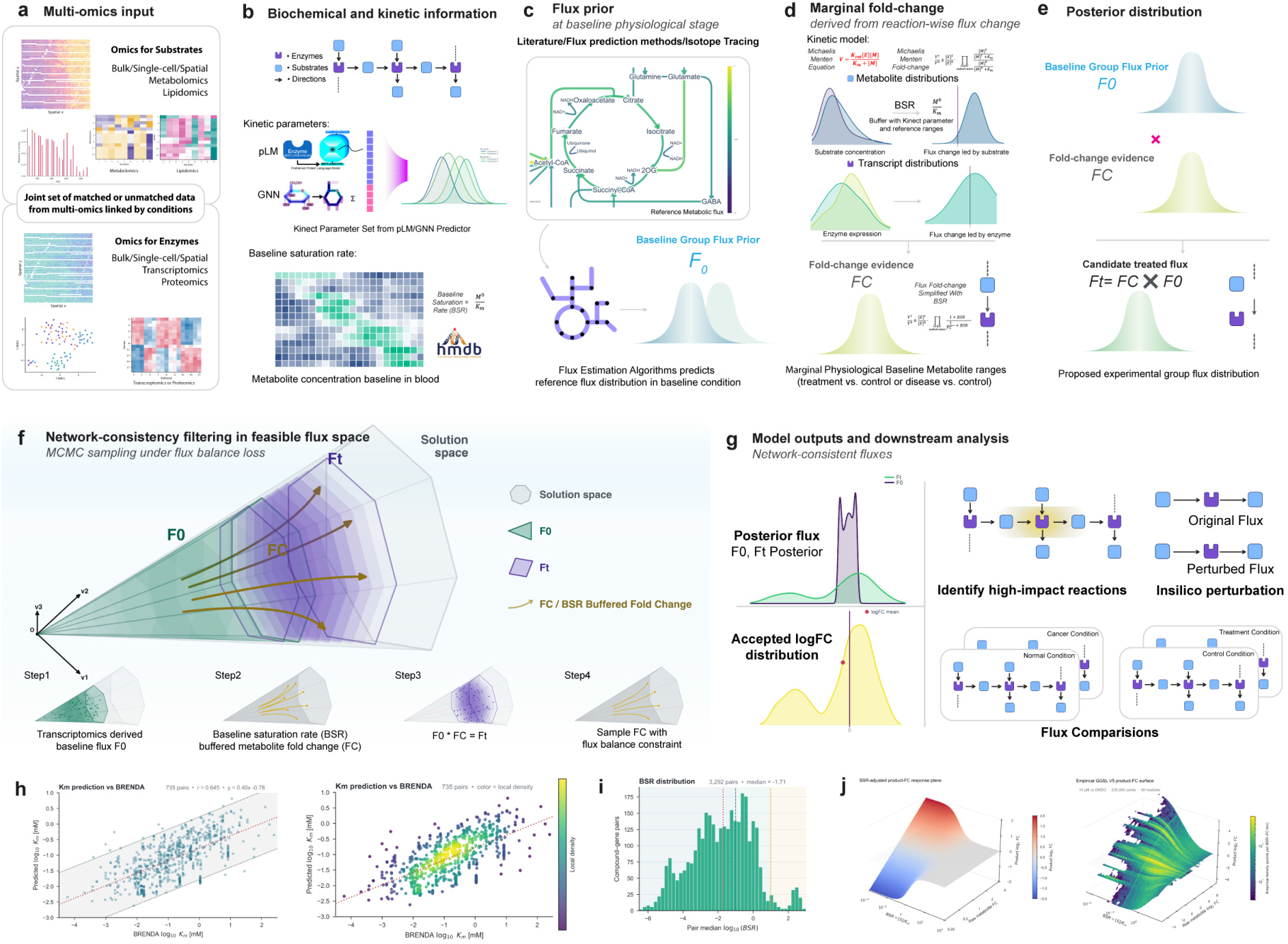
Overview, mathematical formulation, and kinetic validation of mmFEA. **(a)** mmFEA accepts substrate-associated measurements from bulk, single-cell, or spatial metabolomics and lipidomics, together with enzyme-associated measurements from bulk, single-cell, or spatial transcriptomics or proteomics. The modalities need not be measured in the same samples, provided that observations can be assigned to corresponding biological conditions. **(b)** The curated metabolic network specifies enzyme-substrate relationships, reaction directionality, and subcellular compartmentalization. Missing K_*m*_ values are inferred using a predictor that combines protein-language-model representations with graph neural networks and is benchmarked against BRENDA. [20] [21] [22] Physiological metabolite concentrations from HMDB are used to construct baseline saturation ratio (BSR) distributions. [23] **(c)** The reference-state flux prior *F*_*0*_ can be supplied by published flux measurements, stable-isotope tracing, or a compatible computational flux-estimation method, such as scFEA. [13] [61 **(d)** For each reaction, mmFEA combines the enzyme-associated fold change FC_*E*_ with the Michaelis – Menten-adjusted substrate component 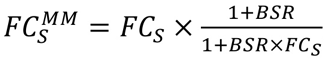 where 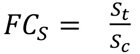 and 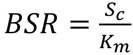, to obtain the marginal reaction-flux fold-change distribution 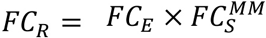. **(e)** Candidate perturbed-state fluxes are obtained by combining reference flux samples with marginal flux-change samples, F_*t*_ F_*0*_ x FC_*R*_, across the network. **(f)** Schematic flux cones illustrate feasible reference fluxes F_*0*_ (green), perturbed fluxes F_*t*_ (purple), and their fold changes *FC*_*R*_ (orange) within the constrained solution space. **(g)** The four-stage workflow estimates *F*_*0*_, constructs *FC*_*R*_, proposes *F*_*t*_, and applies network-level flux-balance constraints to sample jointly feasible perturbed fluxes and flux fold changes. **(h)** Predicted *K*_*m*_ values were evaluated against BRENDA measurements for 735 matched enzyme-substrate pairs represented in the analyzed metabolic network.[22] Scatter and local-density visualizations show log10-transformed values; the Pearson correlation was r = 0.645. The dotted red line denotes the ordinary least-squares fit. **(i)** Distribution of BSR across 3,292 matched compound – gene pairs. For each pair, 3,000 Monte Carlo values were calculated by independently sampling HMDB concentration and GGSL Km parameter set distributions according to 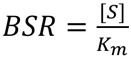.[23] The histogram shows the median log10(BSR) for each pair rather than all Monte Carlo draws. The red dotted line marks the overall median (−1.71); dashed lines delimit BSR regimes below 0.1, from 0.1 to 10, and above 10. (j) Three-dimensional relationship between BSR and metabolite fold change. The left panel shows the theoretical response surface defined by 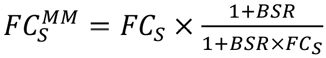. The right panel shows the corresponding empirical surface constructed from 230,000 sampled points across 90 modules of the complete study-specific metabolic network analyzed here, which is not a genome-scale network. The x-, y-, and z-axes represent BSR, raw metabolite log2 fold change, and BSR-adjusted product log2 fold change, respectively. Color represents surface height in the theoretical panel and local data density in the empirical panel. The transparent plane denotes a product log2 fold change of zero.

The computational framework of mmFEA consists of three major steps. First, mmFEA establishes a metabolic flux distribution for the reference state of a selected metabolic network (Figure 1c). Reference distributions can be generated using compatible network-constrained flux estimators. In the analyses reported here, scFEA and MPOCtrL were evaluated as upstream estimators [13] [25]. Experimentally measured reference flux distributions may also be used when available_Jfil. Other computational methods can be incorporated only after their outputs are converted to a compatible reference-flux distribution; Compass was not used as an F_0_ source in the present studyJ11l Consequently, mmFEA is not restricted to a single upstream estimator. Representing the reference state as a distribution preserves uncertainty arising from incomplete data, alternative feasible network states, and variation among reference samples.

Second, mmFEA estimates the marginal distribution of flux changes between the experimental and reference conditions by integrating enzyme- and substrate-associated molecular changes as derived in the formula (4) **(Figure 1d)**. The input of this step includes the reference state flux distribution, enzyme- and substrate-associated molecular changes and the output is a kinetically moderated distribution of flux changes for each reaction. Specifically, transcriptomic or proteomic data are mapped to the enzymes catalyzing each reaction and used to characterize changes in enzymatic capacity. Metabolomic data is mapped to reaction substrates and used to characterize changes in substrate availability. For relative metabolomic inputs, mmFEA uses BSR to account for the unknown absolute scale of metabolite abundance and characterize flux fold change of reactions, while full Michaelis Menton model and pre computed distribution of kinetic parameters are used for quantitative metabolomic inputs. For paired multi-omics datasets, mmFEA retains the correspondence among transcriptomic or proteomic and metabolomic measurements from the same samples. For unpaired data, mmFEA estimates their condition-specific distributions and integrates them probabilistically without sample-level matching, which also accommodates unequal sample sizes and enables modeling with missing data or kinetic parameters.

Third, mmFEA combines the reference flux distribution with the estimated reaction-wise flux-change distributions through network-aware Markov chain Monte Carlo sampling **(Figure 1e**). Candidate flux states are evaluated, sampled and accepted according to their consistency with the metabolic-network structure and a quadratic global flux-balance loss **(Figure 1f**, see details in Methods). Whereas the reference state is expected to be approximately flux balanced, the perturbed state is allowed to exhibit bounded deviations from exact steady-state balance. This soft constraint prevents implausible accumulation or depletion across the network without assuming that disease-associated or experimentally perturbed systems remain at a strict steady state. The sampling procedure therefore generates an ensemble of biochemically coherent flux states rather than forcing the data into a single optimal solution.

The output of mmFEA includes distribution of flux under the experimental and reference states and flux fold change of each reaction **(Figure 1g)**, which support several downstream analyses, namely (1) flux comparison analysis to characterize metabolic landscapes and variations between conditions, (2) metabolic importance analysis to identify key reaction steps and features that determines metabolic changes, and (3) in-silico perturbation analysis to simulate the local and distant impact when perturbing enzyme or substrate levels.

Notably, BSR encodes prior knowledge about substrate saturation under the reference condition. To construct BSR distributions, empirical *K*_*m*_ measurements from BRENDA were used where available, and archived ESM1b compound-enzyme predictions were used for enzyme substrate pairs lacking empirical measurements [20] [21] [22]. Figure 1h compares predicted *K*_*m*_ values with experimentally reported *K*_*m*_ measurements from BRENDA [22]. Figure 1i shows BSR distributions computed from *K*_*m*_ distributions and generic physiological concentration priors derived from HMDBm. Figure 1j illustrates how BSR modifies the effect of substrate fold change in synthetic and human metabolomics data (see details in Supplementary Methods). Together, Figures 1h – j show that predicted *K*_*m*_ values are consistent with reported *K*_*m*_ measurements and that the resulting kinetic priors span relevant saturation regimes and support their use in relative flux inference.

### mmFEA accurately predicts metabolic changes in pancreatic cancer cell line system

To evaluate the accuracy and robustness of mmFEA predictions from heterogeneous and partially matched omics data, we established an in-house multimodal benchmark centered on APEX1 perturbation in pancreatic ductal adenocarcinoma (POAG) models **(Figure 2a)**. Pa03C cells [31] grown as three-dimensional spheroids were treated on days 4 and 6 with one of five APEX1 inhibitors—APX2014, APX2009, APX2051, APX2053 or APX2059-or with OMSO and collected on day 7 for bulk RNA sequencing. APX2014, APX2009 and APX2051 were active inhibitors, whereas APX2053 and APX2059 —served as negative controls.[32] For the APX2014 and OMSO conditions, we additionally generated condition-linked metabolomics profiles and mitochondrial functional measurements, enabling direct comparison between computationally inferred flux changes and experimentally observed metabolic activities. To assess robustness beyond a single experiment, we also incorporated an independently generated single-cell RNA sequencing dataset from Pa03C cells following RNAi-mediated APEX1 depletion. [26] [27] A separate Supplementary analysis examined an APX2014-context, metabolism-focused CRISPR/Cas9 knockout screen in MIA PaCa-2 cells for exploratory candidate prioritization rather than validation of the Pa03C flux estimates.[28] The Pa03C multimodal data were used to evaluate predicted metabolic changes, compare mmFEA with single-omic, pathway-scoring and flux-inference baselines, and assess consistency across related experimental contexts; the CRISPR analysis was interpreted separately as exploratory candidate prioritization.

**Figure 2.**
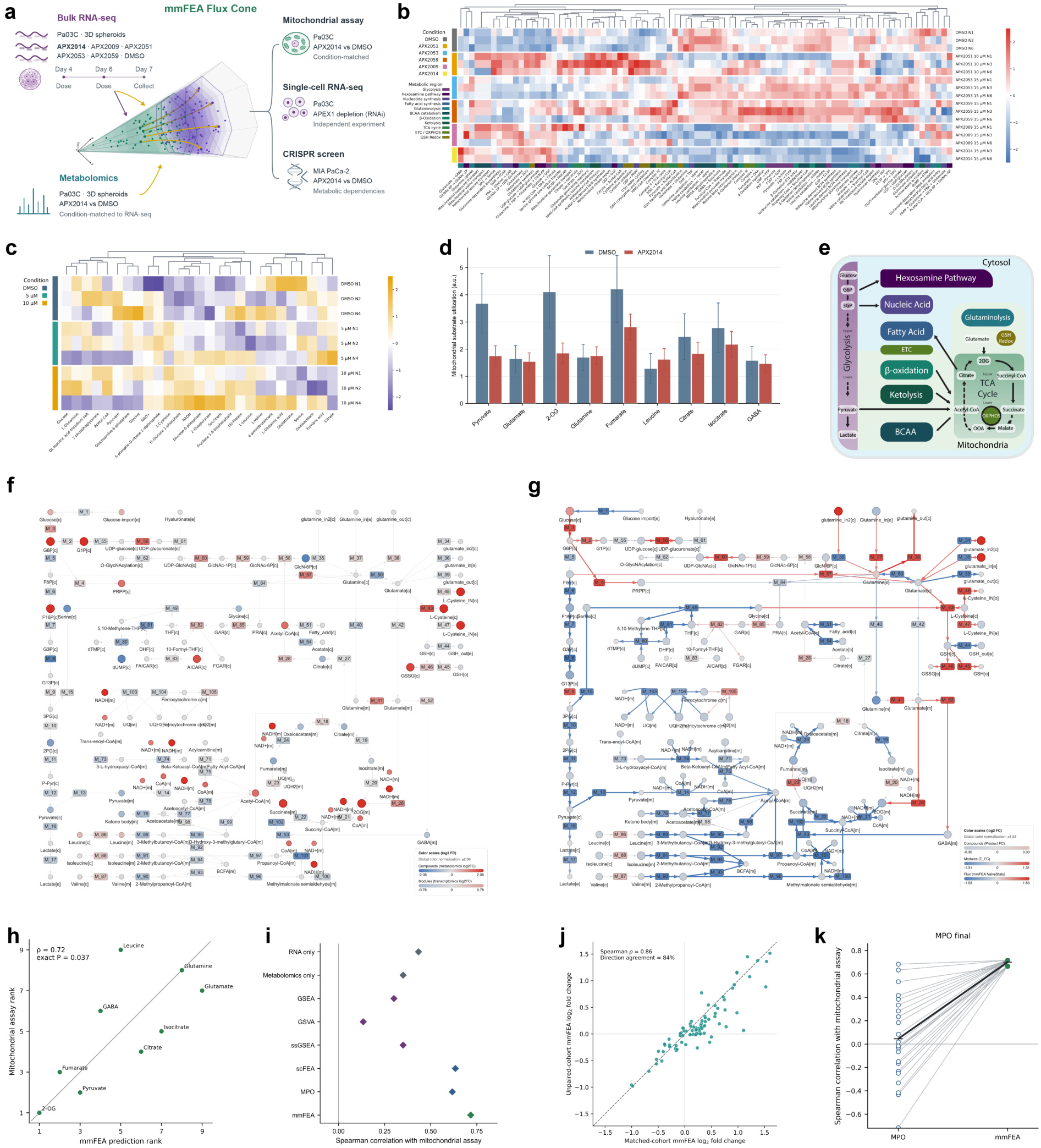
Experiments, metabolic network, and visualization of results. Multimodal evaluation of model-derived relative metabolic-flux changes in APX2014-treated Pa03C cells. **a**, Bulk RNA-seq was generated in replicate across six treatment conditions. For the condition-matched multimodal analysis, we used APX2014-treated and DMSO-control Pa03C spheroids with corresponding metabolomics profiles. Bulk RNA-seq gene-expression data were used to construct F_0_, while metabolomics intensities were incorporated to estimate the marginal FC distributions within the mmFEA pipeline. Predictions were further evaluated using mitochondrial functional assays comparing APX2014-treated and DMSO-control Pa03C spheroids and an independent single-cell RNA-seq dataset following RNAi-mediated APEX1 depletion. [26] [27] An independent metabolism-focused CRISPR screen in MIA PaCa-2 cells was analyzed separately in the Supplementary analysis to prioritize candidate combination targets.[28] In the schematic, green denotes F_0_, purple denotes the predicted treated flux distribution (F_t_), ochre trajectories represent FC, and gray indicates the surrounding feasible flux space. **b**, RNA heat map for the primary GGSL features. Linear CPM values were transformed as log2(CPM + 1), module profiles were calculated as the arithmetic mean of all measured member genes on the log scale, and each displayed feature was standardized across samples. The 50 most variable profiles were selected by across-sample standard deviation without using condition labels. Columns were clustered by average linkage using Euclidean distance; samples were ordered by condition and replicate. The 17 samples comprised DMSO (n = 3), APX2009 (n = 3), APX2014 (n = 2), APX2051 (n = 3), APX2053 (n = 3) and APX2059 (n = 3). Metabolic-region colors denote categorical network assignments, not activity or effect magnitude. **c**, Heat map of 27 quantified network-matched metabolites, including measured malate, across DMSO, 5 µM APX2014 and 10 µM APX2014 samples (n = 3 per condition). Values are normalized log2 MS1 intensities standardized within metabolite across samples; metabolite columns were clustered. **d**, Mitochondrial substrate-utilization measurements for the nine benchmark compounds. Duplicate measurements were averaged within compound, date and condition. Bars show the mean and SEM across five date-matched experiments for DMSO and APX2014. **e**, Functional-region annotation of the 90-module GGSL V5 reduced network. Metabolic regions are color-coded as follows: glycolysis, deep purple; hexosamine pathway, dark plum; nucleic acid synthesis, indigo; fatty acid synthesis, deep blue; glutaminolysis, blue-teal; BCAA catabolism, dark blue-teal; 13-oxidation, jade teal; ketolysis, dark evergreen; TCA cycle, deep green; ETC/OXPHOS, olive green; and GSH redox, ochre. Light cyan and light green denote the cytosolic and mitochondrial compartments, respectively. **f**, Raw-omics network map. Module nodes show the mean member-gene RNA log2 fold change for APX2014 versus DMSO (APX2014 n = 2; DMSO n = 3), and compound nodes show the mean 10 µM APX2014-minus DMSO difference in normalized log2 metabolite intensity (n = 3 per condition). Gray edges show network topology and do not represent inferred flux. The map includes 324 measured network genes and 32 displayed metabolite features (31 mapped compound). **g**, Integrated mmFEA prediction results on GGSL network. Compound nodes encode product-associated fold change, module nodes encode the expression-associated component (E-FC), and edges encode the mean retained log2 flux fold change. The analysis used the 90-module reduced network, 10 µM APX2014-versus-DMSO metabolomics and APX2014-versus-DMSO RNA. **h**, Rank concordance between the primary mmFEA prediction and the mitochondrial assay across nine compounds. The model predicts the log2 change in the metabolic flux consuming each compound. Spearman ρ = 0.717, exhaustive two-sided exact P = 0.0369 and Pearson r = 0.658. i, Comparison of mmFEA with single-omic, pathway-scoring and flux-inference baselines using the same nine compounds. One value is shown for each deterministic method. For scFEA and MPOCtrL (Message Passing Optimization-based Constrained Learning; MPO)[25], the displayed value is the arithmetic mean of the three best benchmark-performing initializations among 25 evaluated initializations. The resulting mean correlations were ρ = 0.633 for scFEA (seeds 15, 19 and 128) and ρ = 0.617 for MPO (seeds 7, 32 and 14). These outcome-selected averages provide optimistic descriptive summaries; no selection-adjusted inference was performed. The primary mmFEA result used T = 0.01 and 9,000 retained post-burn states. **j**, Sensitivity of POAG module-level mmFEA effects to replacement of the transcriptomic cohort and baseline-flux estimate. The matched arm used GSE62452 RNA [29] and its corresponding MPO-derived baseline, whereas the unpaired arm used RNA from the independent CPTAC POAG cohort[30] (135 tumors and 19 adjacent-normal samples) and a CPTAC-derived MPO baseline. The original POAG metabolomics, metabolic network and kinetic inputs were retained in both arms. Across 90 modules, Spearman ρ = 0.862, Pearson r = 0.901 and directional agreement was 84.4%. **k**, Additional initialization stress test evaluating mmFEA under deliberately adverse baseline-flux estimates. Twenty-five MPO-final initializations generated a broad range of F_0_ solutions, including poorly performing, low-agreement initializations intentionally retained to challenge mmFEA robustness rather than characterize typical MPO performance. Lines connect each MPO prediction with its corresponding mmFEA result. Agreement with the mitochondrial assay improved after mmFEA for all 25 initializations. Computational initializations were not treated as biological replicates or inferential degrees of freedom.

Projection of gene expression onto the reaction-module space of the subcellularly resolved GGSL network indicated increased glutamate metabolism, de novo glutathione synthesis, upper glycolysis and branched-chain amino-acid (BCAA) transport, together with decreased tricarboxylic acid (TCA) cycle activity, electron transport, glycosaminoglycan synthesis, lower glycolysis and BCAA catabolism **(Figure 2b**). Metabolomics showed less consistent changes among glycolytic and TCA-cycle intermediates **(Figure 2c)**, whereas the mitochondrial assay indicated decreased utilization of TCA-cycle substrates and moderately increased BCAA associated activity **(Figure 2d). Figure 2e** summarizes the functional regions and reaction modules of the GGSL network used for the raw-omics and mmFEA maps in **Figure 2f-g**. Direct mapping of transcriptomic and metabolomic changes revealed disagreement at several reaction steps, complicating interpretation of either modality alone **(Figure 2f)**. Integration with mmFEA produced a coherent set of predicted changes, including decreased glycolysis, TCA-cycle activity and BCAA catabolism and increased glutamate metabolism and glutathione synthesis **(Figure 2g)**. The predicted compound-consuming flux changes were concordant with the ordering of mitochondrial substrate-utilization responses **(Figure 2h)**. In the cross-method comparison, mmFEA was evaluated with single-omic, pathway-scoring and flux-inference baselines using the same nine compounds. **Figure 2i** displays the arithmetic mean of the three best benchmark performing initializations for scFEA and MPO. Replacing the matched RNA cohort and corresponding baseline F_0_ with an independent CPTAC PDAC cohort preserved the overall module-level response **(Figure 2j). Figure 2k** presents a separate adverse-initialization stress test using 25 MPO-final solutions spanning poorly to strongly agreeing upstream estimates. Agreement with the mitochondrial assay improved after mmFEA for all 25 MPO-final initializations. A parallel scFEA initialization analysis was retained as a supplementary sensitivity analysis.

## DISCUSSIONS

We developed mmFEA as a kinetic-aware statistical learning framework for estimating condition-specific metabolic flux changes from multi-omics measurements when direct fluxomic data are unavailable. The central advance is to connect changes in enzyme-associated measurements and metabolite substrate abundance through Michaelis-Menten kinetics rather than treating transcriptomic or metabolomic signals as direct surrogates of flux. By introducing BSR, mmFEA explicitly models how the same change in substrate abundance can have different effects on reaction rate depending on the baseline kinetic state. The framework further represents molecular measurements, kinetic parameters, and reference fluxes as distributions and uses network-level sampling to identify biochemically coherent flux states while allowing perturbed systems to deviate from strict steady-state assumptions. Importantly, mmFEA estimates relative, model-based flux changes rather than absolute metabolic fluxes, positioning it as a complement to isotope-tracing-based metabolic flux analysis rather than a replacement for direct measurement.

The APX2014 perturbation experiments illustrate the value of this multimodal integration. Transcriptomic and metabolomic measurements alone showed partially discordant metabolic responses, particularly across glycolysis and the TCA cycle, whereas mmFEA yielded a more coherent pattern characterized by decreased glycolysis, TCA-cycle activity, and BCAA metabolism together with increased glutamate metabolism and glutathione synthesis. The biological pattern is consistent with substantial remodeling of mitochondrial and redox metabolism following APEX1 inhibition [26], with suppression of central oxidative metabolismaccompanied by increased glutamate utilization and glutathione production. More generally, this example illustrates why enzyme expression, or metabolite abundance alone may be insufficient for interpreting metabolic adaptation.

A practical strength of mmFEA is that it was designed around the data structures commonly available in human disease studies, where multi-omic measurements are often incomplete and not collected fom matched samples, rather than requiring fully matched multi-omic samples. Because the enzyme- and substrate-associated terms are separable in the reaction-rate fold change formulation, condition-linked transcriptomic and metabolomic datasets can be integrated probabilistically even when generated from different cohorts. This property is supported by the PDAC sensitivity analysis, in which replacing the matched RNA cohort and corresponding baseline-flux estimate with an independent CPTAC cohort produced highly concordant module level effects. The pan-cancer and spatial analyses further suggest that mmFEA can characterize context-dependent metabolic variation across diseases, tissue sites, and spatially resolved samples where direct flux measurements are difficult to obtain. flexibility may be particularly useful for retrospective clinical cohorts, archived specimens, and emerging spatial multi-omic studies.

Several limitations define the appropriate interpretation of mmFEA and motivate future development. Its predictions depend on the quality of the reference-flux distribution, enzyme abundance proxies, metabolic-network representation, and available kinetic information, while baseline metabolite concentrations are currently based on generic physiological priors rather than direct intracellular measurements in the modeled system. In addition, the retained Monte Carlo states do not constitute a formally calibrated Bayesian posterior or biological replicates; their variation reflects uncertainty induced by the specified inputs and computational procedure. Future work should evaluate mmFEA against larger isotope-tracing datasets, improve kinetic and compartment-specific modeling, and incorporate additional regulatory layers such as protein abundance and post-translational modification to strengthen its utility for identifying metabolic mechanisms and vulnerabilities.

## METHODS

### Study design and analysis scope

mmFEA was evaluated in four settings: a Pa03C perturbation experiment with bulk transcriptomic, metabolomic and mitochondrial functional measurements; a tumor–normal pan cancer collection; an unpaired-cohort pancreatic ductal adenocarcinoma (PDAC) sensitivity analysis; and a spatial transcriptomic-metabolomic dataset. Each biological contrast and network version was analyzed separately. Results were not averaged across cancer types or network versions.

The analysis comprised four stages. Transcriptomic data were used as linear CPM values for mmFEA; log_2_(CPM + 1) values were used only for visualization and enrichment analyses. Metabolomic MS1 intensities were normalized and then log_2_-transformed. Expression and metabolite changes were subsequently mapped to metabolic modules and combined with kinetic priors to construct module-level fold-change distributions. These distributions were applied to a baseline flux distribution, *F*_0_, to propose perturbed states, *F*_t_. Proposed states were accepted or rejected according to a normalized flux-balance loss. The resulting quantities are relative, model-based flux changes; they are not absolute metabolic fluxes.

### Pa03C transcriptomic matrix

Analysis began from a supplied linear CPM-scale expression matrix containing 17 Pa03C samples: DMSO *(n* = 3), 15 µM APX2009 *(n* = 3), 15 µM APX2014 *(n* = 2), 10 µM APX2051 *(n* = 3), 15 µM APX2053 *(n* = 3) and 15 µM APX2059 *(n* = 3). Concentrations, compounds and replicate identifiers were parsed from the matrix column labels. APX2053 and APX2059 were negative-control compounds. The primary mmFEA comparison used APX2014-treated samples and DMSO controls, and the linear values were supplied to mmFEA using the linear-input setting. For visualization and enrichment analyses, values were transformed as log_2_(CPM + 1). Module profiles were calculated as the arithmetic mean of all measured member-gene values on this log scale. Heat-map features were selected without reference to treatment labels, using across-sample standard deviation, and were standardized within feature across samples. These standardized values were used only for visualization.

### Pa03C metabolomics

Analysis began from a supplied aligned HILIC negative-ion MS1 feature-intensity matrix containing nine biological samples (DMSO, 5 µM APX2014 and 10 µM APX2014 in each of three batches), one composite MS/MS acquisition and three blank acquisitions. Aligned centroid-feature intensities supplied the quantitative signal; MS/MS assignment and matching fields were retained as annotation evidence rather than used for quantification. The matrix also provided retention time, mass-to-charge ratio, fill rate, signal-to-noise ratio, reference-RT and reference-mass matching, spectral-similarity fields and injection order.

Zero intensities were treated as missing. Feature intensities were normalized by probabilistic quotient normalization using the per-feature median of the included DMSO samples as the reference distribution and were transformed to log_2_ scale.[33] The primary contrast compared 10 µM APX2014 with DMSO across three experimental batches. Each treatment sample was compared only with the DMSO sample acquired in the same biological replicate and analytical batch; the three paired long_2_ fold changes were then summarized across batches. A no-N4 analysis was retained as a sensitivity analysis because batch N4 showed an atypical normalization factor.

The aligned table contained 11,357 features, of which 1,374 carried a KEGG annotation. Feature-level quality review used fill rate, signal-to-noise ratio, blank-to-sample behavior and control-sample behavior. Candidate features were separated by retention-time subgroup before representative selection so that chromatographically distinct signals were not automatically collapsed. To avoid treating related ion features as independent metabolites, one representative signal was retained per KEGG compound. When a compound had at least one quality-control passing candidate, failed candidates were excluded from selection. Eligible candidates were ranked by their proximity to the median batch-specific fold-change profile, followed by lower across-batch variance, MS/MS support, feature-quality score, signal-to-noise ratio, fill rate and alignment identifier. This procedure yielded 577 KEGG-level measurements. Of these representatives, 460 were selected from quality-control-passing pools; 117 compounds for which all candidates failed the stated criteria were retained with an explicit quality flag.

The main metabolomics heat map contained the 27 quantified metabolites matched to the reduced network, including measured malate. The supplementary heat map contained all 577 mapped metabolites and did not apply a top-50 filter. A separate descriptive analysis summarized HILIC-detected complex-lipid features from the same negative-mode experiment. These signals are therefore described as HILIC-detected complex lipids, rather than as a comprehensive lipidomics assay. All metabolomic quantities represent relative ion-intensity evidence, not absolute concentrations.

The primary integrated analysis used 15 µM APX2014-versus-DMSO transcriptomic data and 10 µM APX2014-versus-DMSO metabolomic data. The 5 µM metabolomics group was included in descriptive visualizations, including the metabolomics heat map, but was not used as the metabolomic input for the primary mmFEA comparison. The integration therefore evaluates whether the two omics datasets show a concordant APX2014 response across these doses; it is not a calibrated dose–response analysis.

### Mitochondrial functional measurements

Mitochondrial substrate-utilization measurements were obtained using an S-1 MitoPlate assay; the assay format for Pa03C cells has been described previously.[26] Five date-matched APX2014 and DMSO experiments were analyzed. Duplicate measurements were first averaged within compound, date and condition. For each compound and date, the response was calculated as the APX2014 value minus its date-matched DMSO value in the original assay units. Bars display the arithmetic mean across the five matched dates, with the standard error of the mean (SEM). These measurements report mitochondrial substrate-dependent functional activity and were not interpreted as direct measurements of pathway flux.

Nine compounds were matched to model predictions using a fixed consumed-flux definition. Let *S* denote the signed compound-by-module matrix and *S*_in_= max(−S, 0) its nonnegative consumption component. For state *k* ∈ {0, *t*}, total flux consuming each compound was *C*_in,k_ = *S*_ni_*F*_k_. The benchmark prediction was Δ.*C* = log2(Cin,t + ε) − log2(Cin,o+ε),where ε prevents taking the logarithm of zero. Thus, the benchmark used the change in total stoichiometry-weighted consumed flux rather than the value of a single compound-associated module. Spearman rank correlation compared the nine predicted compound-level changes with the nine mean assay differences. Because the two quantities have different physical units, the primary analysis assessed concordance of their ordering rather than calibration of their absolute values. Pearson correlation, direction agreement, pairwise ordering and standardized pattern error were secondary summaries.

The exact two-sided Spearman *P* value was obtained by exhaustively enumerating all 9! permutations of the compound labels. Compound-resampling intervals used 10,000 bootstrap samples and describe sensitivity to the composition of this nine-compound benchmark. They do not add biological replicates. The temperature and consumed-flux summary rule were selected during exploratory evaluation against the same benchmark; performance estimates using these settings are therefore internally calibrated rather than an independent external validation. No exact P values or confidence intervals were assigned to the outcome-selected top-three mean scFEA and MPO summaries in Figure 2i because initialization ranking used the same nine compound outcome.

Succinate was excluded before method comparison because its mitochondrial-assay measurements did not pass the study-level data-quality review for a reproducible between-condition comparison. The resulting nine-compound set was fixed and applied identically to every method; no method-specific compound filtering or sign reversal was permitted.

### Comparator methods

RNA-only, metabolomics-only, GSEA [34], GSVA [35], single-sample GSEA (ssGSEA)fl.§1, scFEA[13] and MPO[25] were evaluated against the same locked nine-compound functional benchmark. Assay compounds were matched across data types by KEGG compound identifier. For mmFEA, scFEA and MPO, module outputs were converted to compound-level predictions using the signed GGSL stoichiometric matrix: *S*_in_= max(−*S*, 0), *C*_in_,_*k*_ = *S*_in_*F*_k_, and Δ.*C* = log_2_(Cin,t + ε) −log_2_(*C*_in_,_0_+ ε). Thus, each prediction represented the change in total stoichiometry-weighted flux consuming the assayed compound, rather than a selected single module.

RNA-only and enrichment comparators used a prespecified, outcome-blind KEGG gateway mapping because their scores are gene-set activities rather than network fluxes. Each assay compound was assigned by KEGG identifier to its immediate mitochondrial-utilization reaction or reactions-for example, pyruvate to pyruvate dehydrogenase and pyruvate carboxylase, glutamine to glutaminase, y-aminobutyric acid to 4-aminobutyrate aminotransferase, 2-oxoglutarate to 2-oxoglutarate dehydrogenase and fumarate to fumarase. Reaction genes were obtained from KEGG [38] and restricted to MitoCarta3.0.[38] When more than one gateway reaction was retained, its score was multiplied by the compound stoichiometric coefficient and the weighted scores were summed, not averaged. Because the selected KEGG reactions are annotated as reversible, these compound scores represent substrate-linked reaction activity and should not be interpreted as oriented net consumed flux. A broader mapping to every directly connected KEGG reaction and analyses based on GGSL gene sets were retained as sensitivity analyses and were not pooled with the primary independent KEGG-gateway comparison.

RNA-only, GSEA, GSVA and ssGSEA received the same log2(CPM + 1) matrix and the same batch-adjusted APX2014-versus-DMSO linear model(~ batch+ treatment).[39] The RNA-only score for each gateway reaction was the mean fitted long_2_ fold change of its mapped genes. For GSEA, genes were ranked by the moderated treatment *t* statistic and the normalized enrichment score of each gateway-reaction gene set was retained. [34] For GSVA and ssGSEA, gateway-reaction scores were calculated per sample [35] [36] and the treatment coefficient from the same linear model was retained. These reaction scores were projected to assay compounds using the fixed stoichiometry-weighted sum described above. They are expression-derived, dimensionless activity scores and were not relabeled as 1092 flux fold changes.

The metabolomics-only comparator was the within-batch-paired 10 µM APX2014-versus-DMSO log2 abundance fold change of the corresponding KEGG-matched assay compound, averaged only when multiple retained rows shared the same identifier; it was not transformed into a flux value. scFEA and MPO received the same frozen linear-CPM matrix, although their internal transformations remained method specific. Because the resulting scales were not interchangeable, Spearman rank correlation with the mitochondrial assay was the primary cross-method statistic. For scFEA and MPO, Figure 2i displays the arithmetic mean of the three best benchmark-performing initialization-specific correlations among 25 evaluated initializations. These outcome-selected averages provide optimistic descriptive summaries, and no selection-adjusted inference was performed. Figure 2k separately presents an adverse-initialization stress test using 25 MPO-final solutions spanning a broad range of upstream agreement, including poorly performing solutions intentionally retained to evaluate the sensitivity of mmFEA to imperfect F_0_ estimates. Computational initializations were not treated as biological replicates or inferential degrees of freedom. Comparisons used the 90-module reduced network; results from the larger network and complex-lipid analysis were not pooled into the primary benchmark.

### Pan-cancer and unpaired PDAC analyses

The pan-cancer analysis used the Cancer Atlas of Metabolic Profiles. [40] [41] Samples were linked through the supplied MetabID, RNAID and CommonID identifiers, and tumor status was taken from the accompanying manifest. Primary comparisons were restricted to cohorts containing at least 10 tumor and 10 normal samples: BRCA1 (61 tumor, 47 normal), COAD (37, 39), PDAC (27, 12), PRAD (91, 46), ccRCC3 (67, 47) and ccRCC4 (52, 24). GBM (74, 6) and Hi..irthle-cell carcinoma (28, 3) were considered exploratory because of their small normal groups. Transcriptomic transformations followed the source platform: TPM-like matrices remained on a linear scale for mmFEA, whereas log-transformed microarray values were treated as I092-scale expression. Effects were estimated separately within cohort. Processed, non-imputed metabolomics matrices were normalized within dataset, transformed to 1092 scale, filtered for missingness and mapped by KEGG identifier.

The unpaired-cohort PDAC sensitivity analysis compared a matched arm with an arm in which the RNA cohort and its corresponding baseline-flux estimate were replaced. The matched arm used GSE62452 RNA [29] [42], the original PDAC metabolomics data and the matching MPO derived *Fo*. The unpaired arm used CPTAC PDAC RNA from an independent CPTAC PDAC cohort [30] and its independently derived MPO *Fo*, while retaining the original metabolomics component, network and kinetic inputs. After exclusion of adenosquamous specimens, the CPTAC comparison comprised 135 tumors and 19 adjacent-normal tissues. Expression was transformed as log2(value + 1). Integration was performed at the group-contrast level; no sample-level pairing was assumed.

### Spatial transcriptomic-metabolomic analysis

The spatial transcriptomic-metabolomic dataset was obtained from a published spatial multi omic study. [43] [44] Section B1 contained 2,347 locations shared across the required input matrices. Transcript counts were transformed as log(1 + *x*), and MSI intensities remained on a linear scale. For each target location, an empirical local *Fo* distribution was constructed from 8-20 adaptively selected neighboring locations after excluding the target location itself. Spatial locations were regarded as correlated observations; where inference was performed, spatially constrained permutation or block resampling was used rather than treating locations as independent biological replicates.

### Metabolic networks

The 90-module GGSL V5 reduced network was the primary network. It was assembled from reaction annotations in KEGG [37], Reactome [45] and Human1[46] and manually curated using published biochemical literature to refine reaction membership, directionality, gene assignments and subcellular localization. The network covers glycolysis, the hexosamine pathway, nucleotide synthesis, fatty acid synthesis, glutaminolysis, branched-chain amino-acid catabolism, 13-oxidation, ketolysis, the tricarboxylic acid cycle, the electron-transport chain and oxidative phosphorylation, and glutathione-associated redox metabolism. Cytosolic and mitochondrial localization was recorded separately from these metabolic-region assignments. The signed compound-by-module matrix, S (implemented as cmMat), records whether a compound is consumed or produced by each module and supplies the balance constraints used by mmFEA. Module gene sets provide expression evidence but do not constitute complete Boolean gene-protein-reaction rules. The reduced network contained 65 compound-balance rows and 90 metabolic modules.

### Kinetic transformation and module fold-change distributions

The kinetic transformation followed Equations 1-4 in Mathematical Considerations and used the same notation: *FC*_*R*_ for the reaction-level fold change, *FC*_*E*_ for the expression-associated component, *FC*_*s*_ for the treatment-to-reference substrate-abundance ratio and FCs^MM^ for its Michaelis-Menten transformation. The dimensionless baseline saturation ratio was *BSR*_*R*_,_*s*_ = [*S*]_*c*_*/ K*_*m*_,_*R*_,_*s*_. These relationships are not rederived here.

The GGSL *K*_*m*_ parameter set assigned evidence in the following order: empirical BRENDA measurement, archived ESM1b compound-enzyme prediction [20] [21], enzyme-level fallback and legacy fallback. Baseline concentrations were taken from the archived HMDB derived concentration resource and were used as generic priors rather than measurements of Pa03C intracellular concentrations. The evidence source and coverage status were retained for every compound-enzyme pair. Construction, fallback logic and uncertainty modeling for the GGSL *K*_*m*_ set are specified in Supplementary Methods.

For a multigene module, treatment and reference samples were concatenated, missing values were replaced by the within-gene median, linear expression was transformed as log(1 + *x*), and each gene was centered across samples. Singular-value decomposition was applied to the resulting matrix. The first right singular vector defined an expression score for each sample and was oriented to agree with the mean module-wide expression shift. Resampled treatment reference score differences generated the expression-associated fold-change distribution. Single-gene modules used resampled treatment-to-reference expression ratios.

Metabolite-associated distributions were generated by sampling treatment and reference abundance, *K*_*m*_ and baseline concentration and applying Equation 3. Expression-associated *FC*_*E*_ and substrate-associated FC_s_^MM^ draws were sampled independently and multiplied according to Equation 4 to obtain *FC*_*R*_. One thousand expression draws and 1,000 combined reaction-fold-change draws were generated per module. Direct, fallback-supported, graph-filled and neutral assignments were recorded separately.

### Baseline flux and balance-based sampling

Baseline-flux distributions were constructed module by module. A single empirical *Fo* value defined a point distribution; two values defined a uniform distribution over their observed range; and three or more values defined a normal distribution using the empirical mean and sample standard deviation, clipped to the observed range. A proposed perturbed state was calculated componentwise as

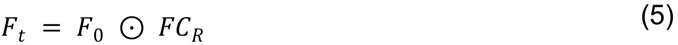

where ⊙ denotes elementwise multiplication. A proposed state was evaluated using the normalized balance loss

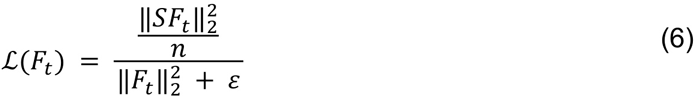

where *S* is the signed compound-by-module matrix, *n* is the number of modules and ε = 10-^12^ prevents division by zero. The loss is a scale-normalized quadratic imbalance: it penalizes imbalance relative to total flux magnitude and prevents the trivial solution obtained by shrinking every flux toward zero. It does not impose exact steady-state balance, and division by *n* is a fixed scaling convention rather than an inferential degree-of-freedom correction. For a proposed state 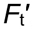*’*, the loss increment and acceptance probability were

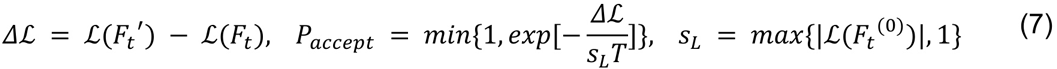

where Tis the sampling temperature and *S*_*L*_ scales the loss by its magnitude at initialization. Proposals that decreased the loss were always accepted; proposals that increased it could be accepted according to Equation 7. In the audited Pa03C runs, the initial normalized loss was less than 1, so the implemented floor gave *S*_*L*_ = 1 and *T* directly controlled the strength of loss based reweighting.

Every proposal independently redrew the complete vectors of *F*_*o*_ and *FC*_*R*_ from their empirical proposal distribution, denoted *q*. Under this independence proposal, the energy-only acceptance rule can be interpreted as sampling the empirical proposal distribution after stoichiometric reweighting:

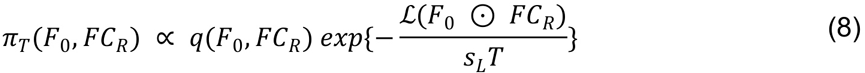

This interpretation explains why an explicit proposal-density ratio is not required: the proposal distribution is also the base measure in Equation 8 and its terms cancel in the acceptance ratio. [47] [48] The resulting distribution is nevertheless not a calibrated Bayesian posterior. The balance loss is a regularization energy rather than a data likelihood, and the temperature was selected empirically rather than assigned a probabilistic calibration. Accordingly, retained-state variation quantifies uncertainty induced by the specified empirical inputs and computational procedure, not biological-population uncertainty.

Iteration zero was stored separately and did not enter the accepted-state, all-state or loss summaries. Following a rejected proposal, the current state was repeated in the retained Markov chain, preserving the residence time required by the reweighted-chain interpretation. The primary Pa03C runs used 10,000 iterations, a burn-in of 1,000 iterations, thinning of 1 and *T* = 0.01, yielding 9,000 retained states. These states are autocorrelated and must not be counted as 9,000 independent observations. Module effects were summarized as the mean retained long2 fold change, which is equivalent to the 1092 of the geometric mean fold change. Retained-state quantiles characterize computational variation under the specified inputs and sampling procedure; they are not confidence intervals for biological-population effects.

Temperature was selected during exploratory analysis against the same nine-compound mitochondrial benchmark and was therefore treated as a tuned computational setting. Sensitivity runs were retained separately. At T = 0.01 in the reduced-network MPO-final lineage, the mean post-burn acceptance rate was 73.8%, and retained module fold changes remained highly correlated with the unfiltered proposal summaries (Spearman ρ = 0.9967). Thus, the selected setting produced moderate stoichiometric reweighting; the shared multi-omic fold change distributions remained the dominant determinant of module ordering. This observation does not imply that F_0_ or the loss filter is irrelevant, because both changed acceptance, retained loss, absolute flux values and some module-level effects. The run length, burn-in, thinning, full vector proposal and loss-scale floor were operational settings rather than parameters established by a formal convergence or efficiency analysis; interpretation therefore relies on the recorded temperature, acceptance, retained-loss and initialization-sensitivity diagnostics. In the principal analysis using the GGSL Km set, the historical product mapping and expression fold change inputs were held fixed while the kinetic resource and Markov-chain outputs were rebuilt. Seed identifiers were algorithm specific: equal integer labels did not define matched scFEA and MPO observations. Outcome-selected top-three initialization averages were used in the cross method display, and the expanded initialization analyses were reported separately.

### Baseline-flux sensitivity

Dependence on the upstream F_0_ estimate was assessed using 25 MPO-final initializations (seeds 1-21, 32, 64, 128 and 256). The expression matrix, metabolic network and MPO settings were held fixed while the initialization seed was varied. The expanded seed set intentionally produced a broad range of F_0_ estimates, including low-agreement solutions that were retained to stress-test mmFEA under imperfect baseline-flux estimation. Each MPO result was propagated through mmFEA using the same reduced network, 10 μ M metabolomics fold change input, T = 0.01 and 9,000 retained post-burn states. The downstream mmFEA random-number stream was held constant within each before-after comparison. These initializations quantify computational sensitivity and were not treated as biological replicates or inferential degrees of freedom. Results were summarized by the distribution of compound-level Spearman correlations before and after mmFEA and by the number of initializations for which agreement increased.

### Statistical analysis and reporting

The experimental unit was defined before analysis: individual samples for the transcriptomic comparison, matched batches for the metabolomics comparison, matched experimental dates for the mitochondrial assay, and subjects within cohort for tumor − normal analyses. Compounds constituted the observations in the mitochondrial concordance benchmark. Monte Carlo draws, *K*_*m*_ values, metabolic modules, retained chain states and computational initializations did not contribute biological degrees of freedom.

The APX2014 RNA comparison included two treated and three DMSO samples and was interpreted as a small-sample analysis. The primary metabolomics contrast included three matched batches. With only three pairs, attainable permutation resolution is limited; effect sizes and consistency across batches were therefore emphasized. Values generated to approximate input distributions were excluded from all experimental sample counts.

Spearman rank correlation was the primary statistic for comparisons between model predictions and mitochondrial functional measurements because the quantities are measured on different scales. Pearson correlation, directional agreement, pairwise-order agreement and standardized root-mean-square pattern error were secondary descriptive measures. Where multiple hypothesis tests were performed within a declared family, Benjamini — Hochberg correction was applied [49] and the family was reported. Tests were two-sided unless a directional alternative had been specified before analysis.

Pan-cancer effects were estimated separately by cancer type. The matched and unpaired PDAC outputs were compared across the same 90 modules using Spearman correlation, Pearson correlation and directional agreement. Module-bootstrap intervals were reported as descriptive sensitivity summaries because modules share genes, metabolites and balance constraints and therefore do not form independent experimental units.

## Data and code availability

Previously reported datasets are identified by citation and accession in the manuscript. Additional data and analysis code are available from the corresponding authors upon reasonable request.

## Notes

### Competing Interest Statement

The authors have declared no competing interest.

